# Semi-supervised Calibration of Risk with Noisy Event Times (SCORNET) Using Electronic Health Record Data

**DOI:** 10.1101/2021.01.08.425976

**Authors:** Yuri Ahuja, Liang Liang, Selena Huang, Tianxi Cai

**Affiliations:** Harvard School of Public Health; Harvard Medical School; Harvard School of Public Health; Brigham and Women’s Hospital

## Abstract

Leveraging large-scale electronic health record (EHR) data to estimate survival curves for clinical events can enable more powerful risk estimation and comparative effectiveness research. However, use of EHR data is hindered by a lack of direct event times observations. Occurrence times of relevant diagnostic codes or target disease mentions in clinical notes are at best a good approximation of the true disease onset time. On the other hand, extracting precise information on the exact event time requires laborious manual chart review and is sometimes altogether infeasible due to a lack of detailed documentation. Current status labels – binary indicators of phenotype status during follow up – are significantly more efficient and feasible to compile, enabling more precise survival curve estimation given limited resources. Existing survival analysis methods using current status labels focus almost entirely on supervised estimation, and naive incorporation of unlabeled data into these methods may lead to biased results. In this paper we propose Semi-supervised Calibration of Risk with Noisy Event Times (SCORNET), which yields a consistent and efficient survival curve estimator by leveraging a small size of current status labels and a large size of imperfect surrogate features. In addition to providing theoretical justification of SCORNET, we demonstrate in both simulation and real-world EHR settings that SCORNET achieves efficiency akin to the parametric Weibull regression model, while also exhibiting non-parametric flexibility and relatively low empirical bias in a variety of generative settings.

## 1 Introduction

The Electronic Health Record (EHR) has in recent years become an increasingly available source of data for clinical and translational research (Kohane *and others*, 2012; Hripcsak and Albers, 2012; Miotto *and others*, 2016). Comprising heterogeneous clinical encounters including diagnostic and procedural billing codes, lab tests, prescriptions, and free text clinical notes for millions of patients, these rich data offer abundant opportunities for in silico epidemiological analysis. One application that has garnered recent interest is estimation of population disease risk within EHR patient cohorts, which can enable more powerful and precise estimation of real-world disease risks as well as comparative effectiveness analysis of alternative treatment strategies (Hodgkins *and others*, 2017; Dean *and others*, 2003; Liu *and others*, 2018; Panahiazar *and others*, 2015; Steele *and others*, 2018). Several studies have had success estimating time to death within rule-defined disease cohorts (Panahiazar *and others*, 2015; Steele *and others*, 2018). However, estimating the temporal risk of developing a disease is a more challenging task due to EHR’s lack of direct observations of either disease status or the timing of disease onset. Convenient proxies of disease status or onset time based on readily available features such as International Classification of Disease (ICD) codes often exhibit low specificity and systematic temporal biases, potentially yielding highly biased disease risk estimators if used as event time labels (Cipparone *and others*, 2015; Uno *and others*, 2018). On the other hand, extracting precise information on disease outcomes requires labor-intensive manual chart review, which is particularly challenging for event times since the event may occur outside of the hospital system and only be mentioned during follow-up visits. It is thus only practically feasible to annotate the current status Δ = *I* (*T* ≤ *C*) of the event time *T* , where *C* is the follow up time.

In this paper, we consider the problem of estimating the disease risk *F* (*t*) = *P*(*T* ≤ *t*) when only a small number of labels onΔ and a large quantity of unlabeled EHR features **W**, including proxies of *T* , are available. Supervised survival curve estimation with current status data on {Δ, *C*} is well established in the statistical literature with several available parametric, semi-parametric and non-parametric procedures (Vardi, 1982; Huang, 1996; van der Laan and Robins, 1998; van der Laan and Jewell, 2003; Lin *and others*, 2019, e.g.). For example, van der Laan and Robins (1998) proposed a non-parametric, locally efficient estimator via inverse probability of censoring weighting (IPCW), assuming that (1) *T* and *C* are conditionally independent given some informative baseline covariates **Z**_**0**_ ⊂ **W** (e.g. age, sex, etc.) and (2) a consistent estimator for the conditional density of *C* | ***Z***_0_ is available. However, these existing estimators do not leverage unlabeled EHR feature information such as time to first surrogate ICD code, which may greatly improve risk estimation precision.

Since **W** may be highly predictive of *T* , the estimation of *S*(*t*) can potentially be improved via semi-supervised learning (SSL) leveraging both the small set of Δ observations in the labeled set and the EHR features **W** in the unlabeled set. SSL has been shown to significantly mitigate bias and/or improve efficiency for various risk prediction applications (Chai *and others*, 2017; Liang *and others*, 2016; Bair and Tibshirani, 2004; Golub *and others*, 1999). For example, several studies employ semi-parametric models to impute event times in the unlabeled set for subsequent input into an outcome survival model alongside labeled data (Chai *and others*, 2017; Liang *and others*, 2016; Zhao *and others*, 2014; Uno *and others*, 2018; Hassett *and others*, 2017; Chubak *and others*, 2012; Choi *and others*, 2015; Kaji *and others*, 2019; Ruan *and others*, 2019; Ahuja *and others*, 2020*a*). While such imputation approaches may improve efficiency under correct specification of the imputation model, they are subject to significant bias if the imputation model is misspecified. In addition, these existing methods do not allow for use of current status labels for training. Other general augmented inverse probability weighting methods in the missing data literature (Seaman and White, 2013; Rotnitzky and Robins, 2014, e.g.) are not directly applicable here since the probabilities of labels being observed tend to zero in the SSL setting.

We address this shortcoming by proposing Semi-supervised Calibration of Risk with Noisy Event Times (SCORNET) for estimation of *S*(*t*). SCORNET utilizes current status labels while also employing a robust semi-supervised imputation approach on the extensive unlabeled set to maximize survival estimation efficiency. To mitigate imputation bias and maximize efficiency gain from the unlabeled data, SCORNET utilizes a highly flexible semi-non-parametric kernel regression model with EHR features as covariates, which ensures the validity of the resulting risk estimator without requiring the imputation model to hold. In addition to providing theoretical justifications for the SCORNET estimator, we illustrate via simulation studies that SCORNET substantially outperforms existing methods with regards to the bias-variance tradeoff. The rest of the paper is organized as follows. In Section 2, we detail the SCORNET procedure along with its associated inference procedures. In Section 3, we report risk estimation performance relative to existing methods in diverse simulation studies. To further illustrate the utility of SCORNET in clinical applications, we apply it to a real-world EHR study estimating the risk of heart failure among rheumatoid arthritis patients in Section 4. Finally, in Section 5 we briefly discuss the strengths, weaknesses, and potential applications of SCORNET.

## 2 Methods

### 2.1 Setup

Let *T* denote the event time for which we are interested in estimating a cumulative distribution function *F* (*t*) = *P*(*T* ≤ *t*) and survival function *S*(*t*) = 1 − *F* (*t*). In the EHR study we do not observe *T* but rather Δ = *I* (*T* ≤ *C*) for a small labeled subset, where *C* is the follow up time with finite support [0, *τ*_*c*_]. For all subjects, we also observe a set of baseline covariates ***Z***_0_ and longitudinal EHR features ***Z***. Since codes used in the EHR are often highly sensitive but not specific, there often exists some filter variable 𝔽 ∈ {0, 1} such that Δ_*i*_| (F_*i*_ = 0, **W**_*i*_) = 0 almost surely, where 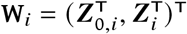. Moreover, we assume that (*T*, ***Z***)*C* | ***Z***_0_. We assume that data for analysis consist of a small set of *n* current-status-labeled observations randomly selected among those with 𝔽 = 1 along with a larger set of *N* unlabeled observations: 𝔻 = {**D**_*ii*_= (*C*_*ii*_, *V*_*ii*_Δ_*i*_, **W**_*i*_, *V*_*i*_, 𝔽_*i*_)^T^,*i* = 1, …, *N*} = 𝕃 ∪ 𝕌, where 𝕃 = {(*C*_*i*_, Δ_*i*_, **W**_***i***_, 1, 1)^T^: 𝔽_*i*_ = 1, *V*_*i*_ = 1, *i* = 1, …, *N*} and 𝕌 = {(*C*_*i*_, 0, **W**_***i***_, 0, 𝔽_*i*_)^T^: *V*_*i*_ = 0, *i* = *n* + 1, …, *N*} with log(*N*)/log(*n*) → *ν*_0_ *>* 3/2 as *n* → ∞.

Since the censoring *C* may depend on ***Z***_0_, we follow the IPCW strategy of van der Laan and Robins (1998) to weight observations by

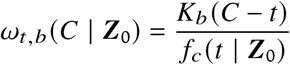

where *Kb*(*s*) = (*s*/*b*)/*b, f*_*c*_ (*t* | ***Z***_0_) = *F*_*c*_ (*t* | ***Z***_0_)/ *t, F*_*c*_ (*t* | ***Z***_0_) = *P*(*C* ≤ *t* | ***Z***_0_),*K*(·) is some symmetric density function, and 0 *<* = *0* (*n*^−*ν*^) is the bandwidth thereof with *ν* ∈ (1/5, 1/3]. IPCW enables consistent estimation of functionals of *T* ≤ *t* and **W** since for any reasonable choice of function *q* (·) and *a*, ∈ {0, 1},

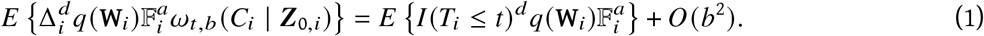

The IPCW estimator for *π*(*t*) = *P*(*T*_*i*_ ≤ *t* | 𝔽_*i*_ = 1) proposed by van der Laan and Robins (1998) essentially corresponds to

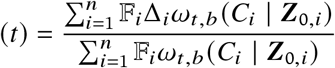

with *f*_*c*_ (*t* | ***Z***_0_) in *ω*_*t,b*_(*C*_i_ | ***Z***_0,*i*_) replaced by a consistent estimator that converges faster than *π*^−1/4^, which is not difficult to achieve under reasonable modeling assumptions since *C*_*i*_ | ***Z***_0,*i*_ can be estimated using the full data 𝔻. To this end, we propose to derive an estimator for the conditional density *f*_*c*_ (*t* | ***Z***_0_) = *λ*_*c*_ (*t* | ***Z***_0,*i*_)*S*_*c*_ (*t* | ***Z***_0,*i*_) by imposing a semi-parametric model for *C* | ***Z***_0_. Although many commonly employed models can be used since once again *C* is fully observed for all patients, we illustrate our proposal by focusing on the Cox proportional hazards model (Cox, 1972) under which

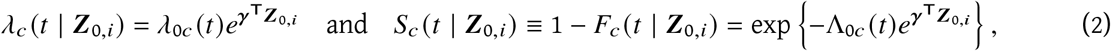

where *λ*_*c*_ (*t* | ***Z***_0,*i*_) is the conditional hazard function of *C*_*i*_| ***Z***_0,*i*_, *λ*_0*c*_ (*t*) is the unknown baseline hazard function, 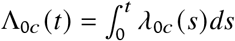, and *γ* is the vector of unknown covariate effects.

### 2.2 SCORNET Estimation

As outlined in Figure 1, SCORNET consists of three steps: (1) estimating the conditional censoring distribution *h* (*t* | ***Z***_0_) using 𝔻; (2) fitting an imputation *working* model for *π*(*t* | **W**) ≡ *P*(*T* ≤ *t* | **W**, 𝔽 = 1) using 𝕃, denoting the estimate of *π*(*t* | **W**) as (*t* | **W**); and (3) estimating *S*(*t*) by marginalizing (*t* | **W**) 𝔽 +Δ (1 − 𝔽) = (*t* | **W**) 𝔽 via IPCW.

**Figure 1:**
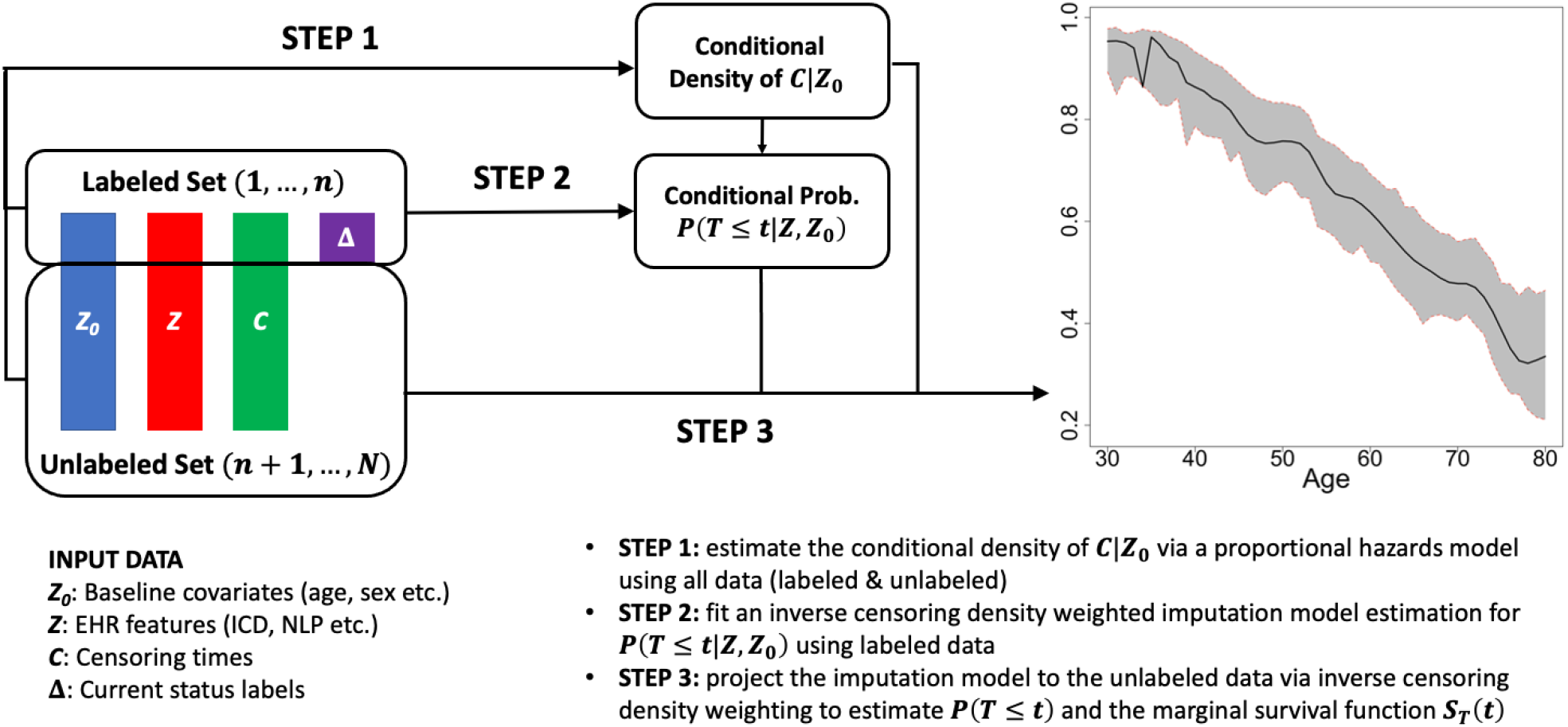
Schematic of the SCORNET algorithm.

#### 2.2.1 Step 1: Estimate *f*_*c*_ (*t* | *Z*_0_) Under the Cox Model for *C* | *Z*_0_

To estimate *f*_*c*_ (*t* | ***Z***_0_), we fit the Cox model (2) to the full data 𝔻 to obtain the partial likelihood estimator 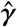 for *γ*. We subsequently estimate Λ_0_ (*t*) and *λ*_0*c*_ (*t*) respectively as the standard Breslow estimator 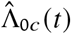 and the kernel-smoothed Breslow estimator 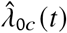 (Basha and Hoxha, 2019), where

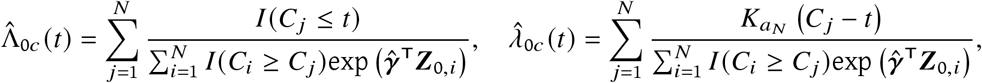

and 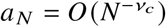 with *ν*_*c*_ ∈ (1/5, 1/3]. We then obtain

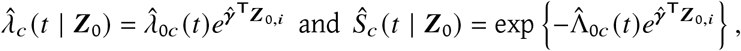

and we estimate *f*_*c*_(*t* | ***Z***_0,*i*_) as 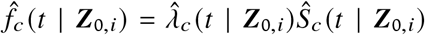. Following standard asymptotic results for non-parametric kernel regression (Pagan and Ullah, 1999), it is not difficult to show that 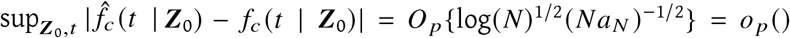. We denote the resulting estimate for the censoring weight as 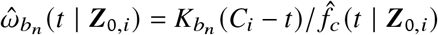.

#### 2.2.2 Step 2: Estimate an Imputation Model *π*(*t* | W_*i*_) ≡ *P* (*T*_*i*_ ≤ *t* | W_*i*_, 𝔽_*i*_ = 1)

To leverage the unlabeled data, we fit a flexible imputation *working* model

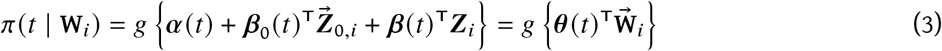

where ***Z***_*i*_ denotes the EHR surrogate features, *θ*(*t*) = (*α* (*t*), *β*_0_(*t*)^T^, *β* (*t*)^T^)^T^, and 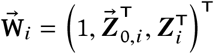. Under (3), 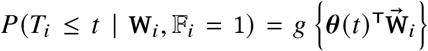, and hence we may estimate ***θ***(*t*) as 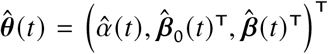, the solution to the IPCW estimating equation evaluated with 𝕃,

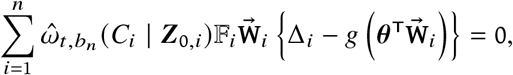

where *b*_*n*_ = *0* (*n*^−*ν*^) with *ν* ∈ (1/5, 1/2). In practice, *b*_*n*_ can be chosen via either standard cross-validation or heuristic plug-in values. For a future observation with filter status 𝔽_*i*_ = 1 and covariates **W**_***i***_, we impute *I* (*T*_*i*_ ≤ *t*) as the conditional risk 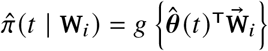.

It is not difficult to show that 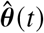 converges in probability to 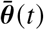, the solution to the limiting estimating equation

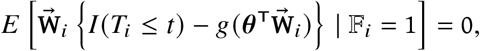

which ensures that

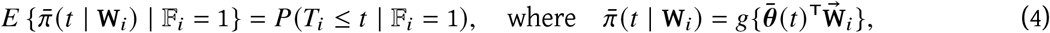

regardless of the adequacy of the imputation model (3).

#### 2.2.3 Step 3: Estimate *F* (*t*) by Marginalizing Imputed Risks

Finally, we marginalize the imputed values 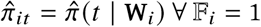 and Δ_*i*_ = 0 ∀ 𝔽_*i*_= 0 to estimate *F* (*t*). Since 𝔽_*i*_ depends on *C*_*i*_, we again employ IPCW to marginalize 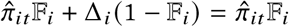 and thereby construct our final estimator for *F* (*t*):

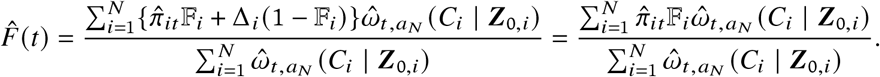

### 2.3 Inference for 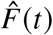

Following standard theory for non-parametric kernel regression (Pagan and Ullah, 1999), we show in the Supplementary Materials that 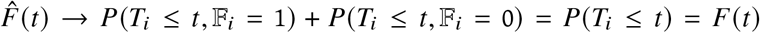 in probability under mild regularity conditions and correct specification of the censoring model regardless of the adequacy of the imputation model. Here, we note that for any *t* ∈ [0, *τ*_*c*_], 0 = *P*(Δ_*i*_ = 0 | 𝔽_*i*_ = 0, *C*_*i*_ = *t*, **W**_***i***_) implies that *P*(*T*_*i*_ ≤ *t* | 𝔽_*i*_ = 0) = 0. Furthermore,

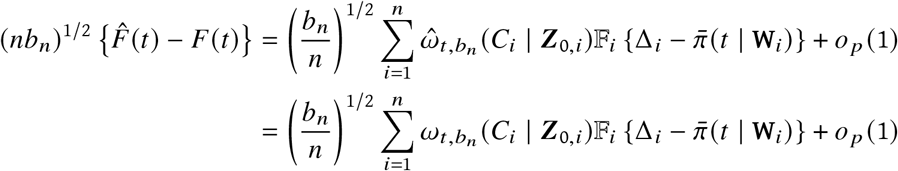

since 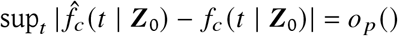. It follows that 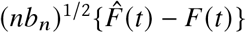 is asymptotically normal with mean 0 and variance

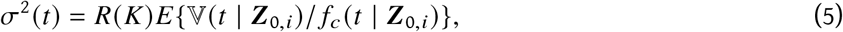

where

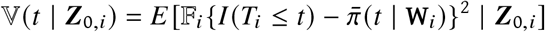

and *R*(*K*) = ∫ *K*(*x*)^2^. Our derivation for the asymptotic distribution of 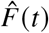 can effectively ignore the variability associated with the estimation of censoring weights, which simplifies the asymptotic variance *σ*^2^ (*t*). Importantly, *σ*^2^ (*t*) decreases as the imputation model approximates *π*(*t* | **W**_***i***_) better since 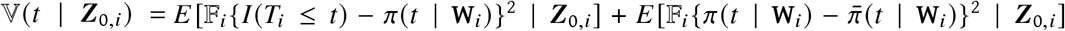 decreases. To estimate *σ*^2^ (*t*) in practice, one may construct a plug-in estimator,

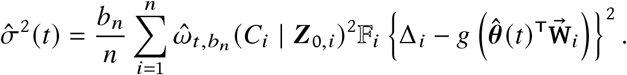

By contrast, the supervised IPCW estimator that incorporates filter negative patients takes the form

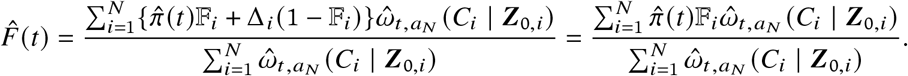

The asymptotic variance of 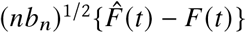 is then *σ*^2^ (*t*) = *R*(*K*)*E* {𝕍 (*t* | ***Z***_0,*i*_)/ *f*_*c*_ (*t* | ***Z***_0,*i*_)}, where

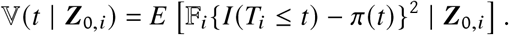

The variance *σ*^2^ (*t*) is equivalent to that of SCORNET if and only if the feature set **W** is uninformative for *T* (i.e. *T* **W**). Supervised IPCW is otherwise less efficient, with relative efficiency controlled by the relative magnitudes of the marginal error *E*[{*I* (*T*_*i*_≤ *t*) − *F* (*t*)}^2^ | 𝔽_*i*_= 1, ***Z***_0,*i*_] and the conditional error 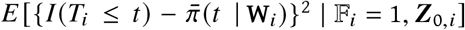.

## 3 Simulation Study

We conduct extensive simulation experimentation to evaluate the finite sample performance of the proposed SCORNET estimator in realistic settings with *n* ∈ {100, 200} observed labels within the set of filter-positive patients, defining the filter to have 99% sensitivity and 88% specificity for Δ. We compare SCORNET to three existing survival function estimators with current status data: 1) parametric Weibull Accelerated Failure Time (AFT) regression with interval event times (Lin *and others*, 2019), 2) semi-parametric Cox Proportional Hazards regression with interval event times and Breslow baseline hazard estimation (Huang, 1996; Cox, 1972; Breslow, 1972), and 3) non-parametric IPCW estimation (van der Laan and Robins, 1998). We incorporate the filter in the Weibull and Cox models by setting | (F = 0) = 0 and weighting the *n* labeled filter-positive patients by 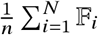. Weibull and Cox are implemented using the icenReg package in R, while IPCW is implemented per the algorithm detailed in van der Laan and Robins (1998), estimating *C* | ***Z***_0_ using 𝔻 under the Cox model. We note that estimating the censoring distribution using 𝕃 yields similar asymptotic performance to using 𝔻, but in finite sample settings the latter offers higher efficiency.

We consider 6 diverse generative mechanisms as detailed in Table 3, including cases where Weibull-distributed accelerated failure time of *T* | ***Z***_0_, proportional hazards of *T* | ***Z***_0_, and proportional hazards of *C* | ***Z***_0_ are respectively violated, as well as cases where SCORNET’s imputation model is and is not misspecified. In settings 1, 2, and 5, we consider various cases where SCORNET and all comparator methods are correctly specified. In setting 1 we consider the specific case where *C* and *T* both depend on ***Z***_0_, and both *C* | ***Z***_0_ and *T* | ***Z***_0_ are Weibull-distributed satisfying accelerated failure time and proportional hazards. In setting 2, by contrast, we consider a case where *T* ***Z***_0_ to assess robustness to over-parametrization of this relationship, and in setting 5 we consider a case where *C****Z***_0_ to evaluate robustness to over-parametrization thereof. In settings 3 and 4 we assess the benefit of SCORNET and IPCW’s robustness to the distribution of *T* | ***Z***_0_ when this distribution satisfies neither Weibull accelerated failure time nor proportional hazards. We evaluate SCORNET’s sensitivity to misspecification of the imputation model in settings 1, 3, and 5, as compared to correct specification thereof in settings 2 and 4. Finally, in setting 6 we assess the sensitivity of SCORNET and IPCW to misspecification of the conditional censoring model *C* | ***Z***_0_. For each given configuration, we compute the empirical bias, standard error, and root mean squared error (RMSE) of all estimators for *F* (*t*) based on their average performance on 500 simulated datasets evaluated at 100 equally-spaced time points *t* ∈ [*Q*_*C*_(0.1) +*b*_*n*_ , *Q*_*C*_ (0.9) − *b*_*n*_], where *Q* denotes the quantile function of *C* under the configuration. We used plug-in bandwidths of = *ŝ* (*C*)*n*^−1/4^ and *a*_*N*_ = ŝ (*C*)*N*^−1/4^ for the imputation (Step 2) and marginalization (Step 3) steps of SCORNET respectively, where ŝ is the empirical standard deviation of observed *C*. We present the performance of the estimators averaged over the selected time points using *n* = 200 labels in Figure 2. The performance at each time point can be found in Supplementary Figure 1, and time-averaged performance using *n* = 100 labels can be found in Supplementary Table 1 of the Supplementary Materials.

**Figure 2:**
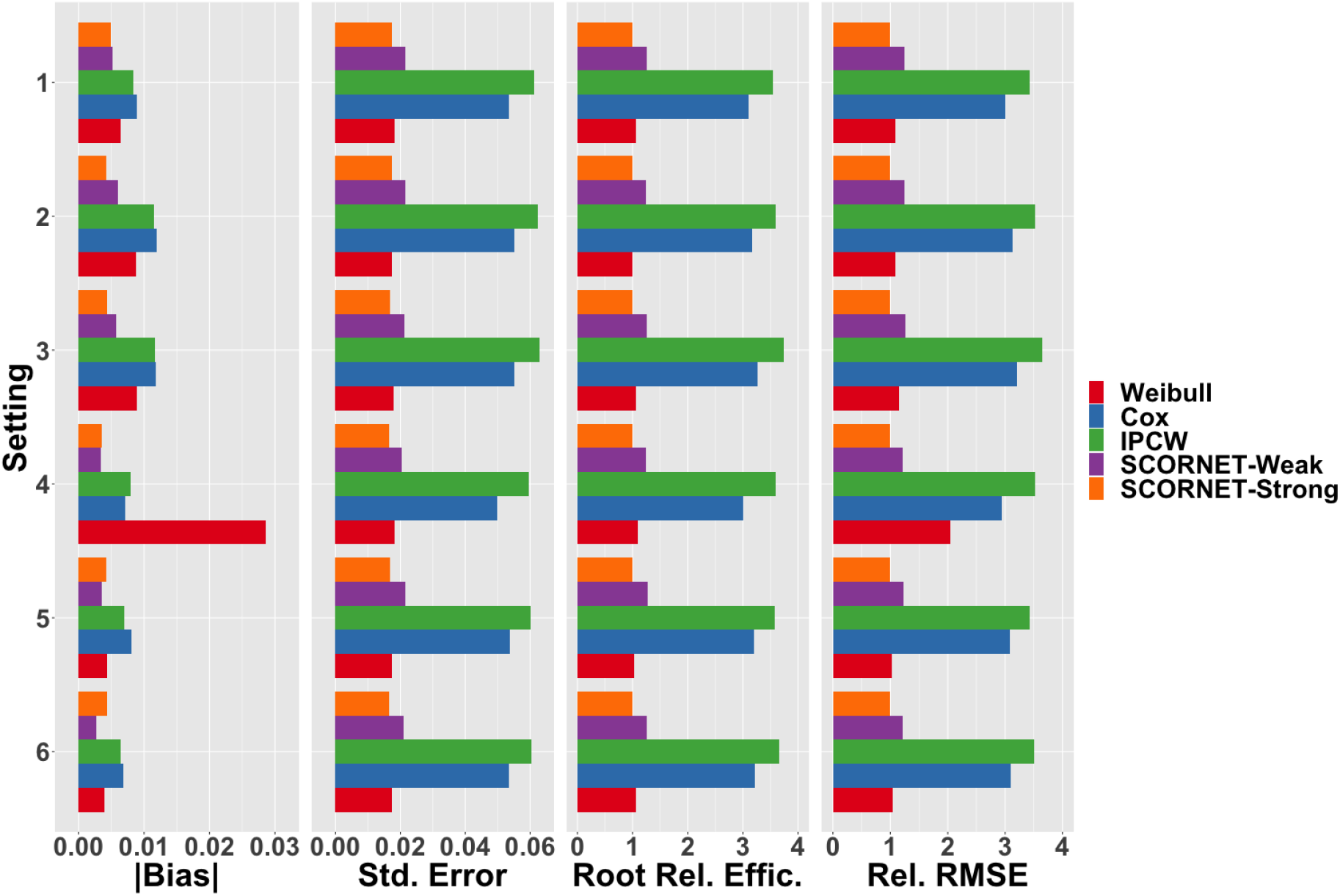
Time-averaged empirical absolute biases (left), standard errors (second from left), root relative efficiencies (second from right), and relative RMSEs (right) of the Weibull Accelerated Failure Time (red), Cox Proportional Hazards w/ Breslow baseline (blue), supervised IPCW (green), and SCORNET estimators using weakly informative (purple) and strongly informative (orange) surrogates, in various simulated settings with *n* = 200 observed current status labels.

As Figure 2 demonstrates, imputing using a strongly informative feature ***Z*** (SCORNET-Strong) results in consistently higher efficiency than just using the weakly informative baseline ***Z***_0_ (SCORNET-Weak), which in turn is markedly more efficient than not leveraging the unlabeled set at all (IPCW). SCORNET makes minimal assumptions regarding the distribution of *T* | ***Z***_0_, settling for non-parametric efficiency in exchange for enhanced flexibility. By contrast, the Weibull regression model fully parametrizes *T* | ***Z***_0_, and the Cox model assumes proportional hazards thereof, increasing efficiency at the expense of bias in the case of misspecification. As expected, Weibull consistently achieves higher empirical efficiency than Cox, which in turn is more efficient than IPCW across settings. Notably, SCORNET consistently achieves empirical efficiency comparable to Weibull and significantly higher than Cox despite being far more flexible than both, again highlighting the efficiency gained by leveraging auxiliary information to impute unobserved risks. At the same time, SCORNET is much less susceptible to model misspecification bias than Weibull, as demonstrated by the latter’s significantly higher bias and RMSE in Setting 4. Indeed, SCORNET achieves relatively low mean absolute bias across settings, with MSE apparently dominated by variance rather than bias in the setting of 100-200 labels. Consistent with the theory, SCORNET is robust to misspecification of the imputation model in settings 1, 3, and 5, achieving equivalently insignificant bias as in setting 2 and marginally but not meaningfully higher bias than in setting 4. That said, correctness of the imputation model in settings 2 and 4 does not yield any meaningful change in relative efficiency, likely because inherent variability functionally dominates imputation model bias given so few labels. Reassuringly, SCORNET (and IPCW) appear insensitive to misspecification of *C* | ***Z***_0_ in setting 6, achieving functionally equivalent bias to the correctly-specified Weibull and Cox models. Altogether, these results corroborate the assertion that SCORNET’s semi-supervised utilization of informative feature data to impute risks in the unlabeled set improves estimation efficiency without introducing bias regardless of the validity of the imputation model. Moreover, they suggest that SCORNET is particularly valuable in settings where (1) flexibility is desired with regard to the distribution of *T* | ***Z***_0_, and (2) there exists a large set of unlabeled patients with associated EHR data – both commonplace in retrospective observational studies.

To assess the finite sample performance of the proposed interval estimation procedures, we obtain standard error estimates both using the proposed plug-in estimator 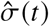 and via bootstrap with 500 replicates. In Figure 3 we demonstrate empirical coverage probabilities of SCORNET’s 95% Wald confidence intervals constructed using each standard error estimator, both averaged over the selected timepoints (left) and at each timepoint (right). Reassuringly, we find that the 95% confidence intervals using both plug-in and bootstrap estimators achieve nearly 95% mean coverage across settings. Coverage only drops below 90% at the tails of the event time support due to moderately increased bias from kernel smoothing thereabout. The plug-in estimator achieves marginally lower coverage than the bootstrap estimator at the right tail due to underestimation of the true standard error, likely because of overfitting of the imputation model given low local censoring density (and thus low effective *N*)Notably, we do not observe this trend in setting 4, wherein correct specification of the imputation model obviates overfitting. Thus, we posit that the plug-in estimator can be reliably used for finite sample problems with *n* ∈ [100, 200] labels as long as the conditional censoring density *f*_*c*_ (*C* | ***Z***_0_) is sufficiently high and the timepoints evaluated are sufficiently far from the tails of the event time support.

**Figure 3:**
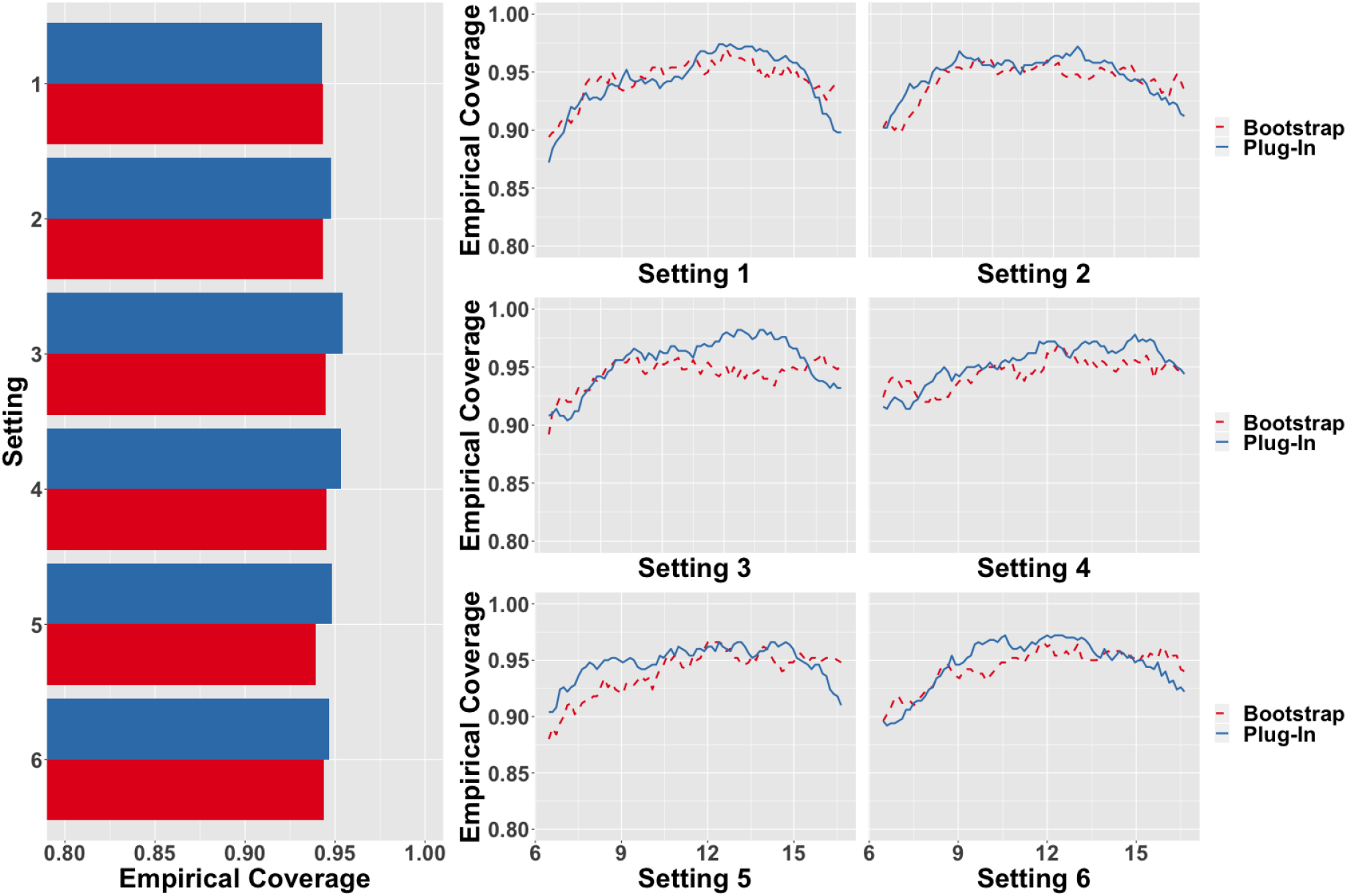
Empirical coverage probabilities averaged over time (left) and plotted over time (right) of SCORNET-Strong’s 95% confidence intervals constructed with the bootstrap (red) and plug-in (blue) standard error estimators in various simulated settings with *n* = 200 observed current status labels. See Table 1 for details of the generative mechanism employed in each setting.

**Table 1:** Generative parameters employed in our simulation study.

| Setting | $\mathbf{Z}_0 \sim$ | $C \mid \mathbf{Z}_0 \sim$ | $T \mid \mathbf{Z}_0 \sim$ | $\mathbf{Z} \sim$ |
| --- | --- | --- | --- | --- |
| 1 | Unif(-1, 1) | Weibull $(10e^{-0.5\mathbf{Z}_0}, \frac{3}{2})$ | Weibull $(15e^{-0.3\mathbf{Z}_0}, \frac{5}{2})$ | Normal$\{T + 1, \sigma(C)/2\}$ |
| 2 | Unif(-1, 1) | Weibull $(10e^{-0.5\mathbf{Z}_0}, \frac{3}{2})$ | Weibull $(15, \frac{5}{2})$ | Normal$\{T + 1, \sigma(C)/2\}$ |
| 3 | Unif(-1, 1) | Weibull $(10e^{-0.5\mathbf{Z}_0}, \frac{3}{2})$ | Weibull $(15e^{-0.3\mathbf{Z}_0^2}, \frac{5}{2})$ | Normal$\{T + 1, \sigma(C)/2\}$ |
| 4 | Unif(-1, 1) | Weibull $(10e^{-0.5\mathbf{Z}_0}, \frac{3}{2})$ | Logistic $(15 - 4\mathbf{Z}_0, 3)$ | Normal$\{T + 1, \sigma(C)/2\}$ |
| 5 | Unif(-1, 1) | Weibull $(10, \frac{3}{2})$ | Weibull $(15e^{-0.3\mathbf{Z}_0}, \frac{5}{2})$ | Normal$\{T + 1, \sigma(C)/2\}$ |
| 6 | Unif(-1, 1) | Weibull $(10e^{-0.5\mathbf{Z}_0^2}, \frac{3}{2})$ | Weibull $(15e^{-0.3\mathbf{Z}_0}, \frac{5}{2})$ | Normal$\{T + 1, \sigma(C)/2\}$ |

## 4 Application to Assessing Heart Failure Risk Among Rheumatoid Arthritis Patients

Rheumatoid arthritis (RA), a chronic inflammatory disease that affects approximately 1% of the general population, is associated with dramatically increased risk of heart failure (HF) morbidity and mortality (Kaplan, 2010; Nicola *and others*, 2005, 2006; Ahlers *and others*, 2020). One study estimated that RA patients have a 1.9-fold lifetime risk of developing HF compared to matched RA-negative controls (Nicola *and others*, 2005), while another estimated that HF accounts for 13% of excess mortality among RA patients (Nicola *and others*, 2006). Ongoing interest lies in estimating the risk of developing HF subtypes in RA cohorts and quantifying the risk modifying effect of various RA treatments (Ahlers *and others*, 2020). Due to the increased availability of electronic health record (EHR) data, it is now possible to assess HF risk for a broader RA population using these data. For example, at Mass General Brigham we previously established an EHR cohort of *N*_0_ = 16, 358 RA patients (Huang *and others*, 2020). This large RA cohort can potentially be used to study the longitudinal risk of HF among RA patients.

However, such analysis is not straightforward as HF status is not readily available within the RA cohort. We propose to estimate HF risk among RA patients by leveraging (1) *n* current status labels on HF status obtained via manual chart review, and (2) informative yet unlabeled EHR data, including time to first ICD code for HF, as surrogate variables ***Z***. We estimate both the age-specific HF risk, *F*_age_ (·), and the risk of developing HF after the patient’s incident ICD code for RA (714), *F*_RA+_ (·), among patients with at least 6 months of follow up whose incident RA codes occur after the age of 16 to select for adult-onset as opposed to juvenile RA. Among filter-positive patients, defined as having at least 1 ICD code for HF, we have *n* = 300 labels on censoring time HF status for age-specific HF risk, and we have *n* = 126 for post-RA HF risk. We let the baseline covariates ***Z***_0_ include sex and decade of first EHR event for *F*_age_ (·), and sex, decade of first RA code, and age at first RA code for *F*_RA+_ (·). We obtain HF risk estimators based on SCORNET as well as the aforementioned comparator estimators. For the imputation model in Step 2 of SCORNET, we consider three EHR-derived surrogate risk predictors for ***Z***: (1) the predicted Δ based on the unsupervised Multimodal Automated Phenotyping (MAP) algorithm, which uses the total counts of HF ICD codes and mentions of HF in clinical notes, as well as the total count of all ICD codes as a healthcare utilization measure (Liao *and others*, 2019), (2) the predicted Δ based on the unsupervised Surrogate-guided Ensemble Latent Dirichlet Allocation (sureLDA) algorithm, which leverages the features used in MAP as well as 121 additional manually-selected EHR features including counts of relevant medications, ICD codes, and concept unique identifiers (CUIs) in clinical notes (Ahuja *and others*, 2020*b*); and (3) the time to first HF ICD code. As in our simulation, we select plug-in bandwidths of *b*_*n*_ = ŝ(*C*)*n*^-1/4^ and *a* _*N*_ = ŝ(*C*) *N*^-1/4^ for the imputation and marginalization steps of SCORNET respectively, and we evaluate risk at 100 timepoints *t* ∈ [*Q*_*c*_(0.1) + *b*_*n*_,*Q*_*c*_(0.9) -*b*_*n*_]. We again compare the performance of SCORNET to that of Weibull, Cox, and IPCW, incorporating the filter in the Weibull and Cox models by propensity weighting as we do in the simulation study.

In Figure 4, we show the estimated HF risk curves along with their standard errors. Reassuringly, all methods appear to agree rather closely for estimation of both age-specific HF risk and HF risk after RA diagnosis. For the latter quantity, however, Weibull and Cox appear to underfit while IPCW appears to overfit relative to the SCORNET estimator, which appears to achieve a reasonable middle ground. Moreover, SCORNET once again attains standard errors comparable to those of the Weibull estimator and meaningfully lower than those of the Cox and IPCW estimators. This suggests that while the Weibull and Cox models potentially fail to capture the complexity of the post-RA HF risk function, and IPCW is too unstable for a limited labeled set of size *n* = 126, SCORNET offers an attractive balance of efficiency and flexibility and is thus well conditioned for such a scenario. As shown in Figure 5, averaged over the selected timepoints, the root relative efficiency of SCORNET is 1.11, 2.55, and 3.31 compared to the Weibull, Cox, and IPCW estimators respectively for estimation of agespecific risk, and 1.34, 2.32, and 3.85 respectively for estimation of HF risk after RA diagnosis. Once again, the fact that SCORNET achieves efficiency moderately higher than the relatively inflexible Weibull model and significantly higher than the Cox and IPCW estimators reflects the value of leveraging available information from the EHR to bolster risk estimation efficiency.

**Figure 4:**
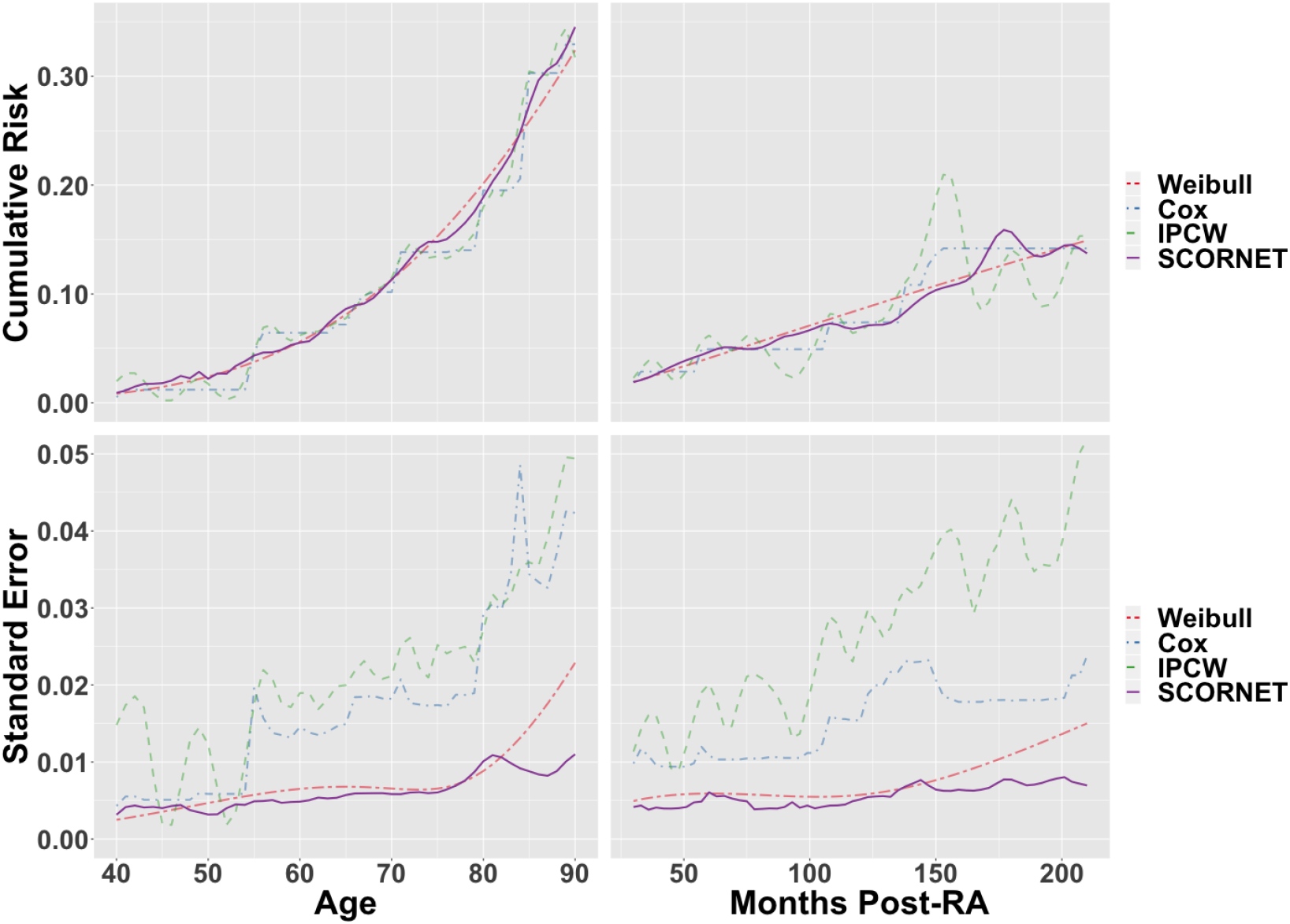
Estimated age-specific and post-RA cumulative risks of heart failure (top) and bootstrap standard errors thereof (bottom) over time of the Weibull Accelerated Failure Time (red, short-long-dashed), Cox Proportional Hazards w/ Breslow baseline (blue, dot-dashed), supervised IPCW (green, dashed), and SCORNET (purple, solid) estimators.

**Figure 5:**
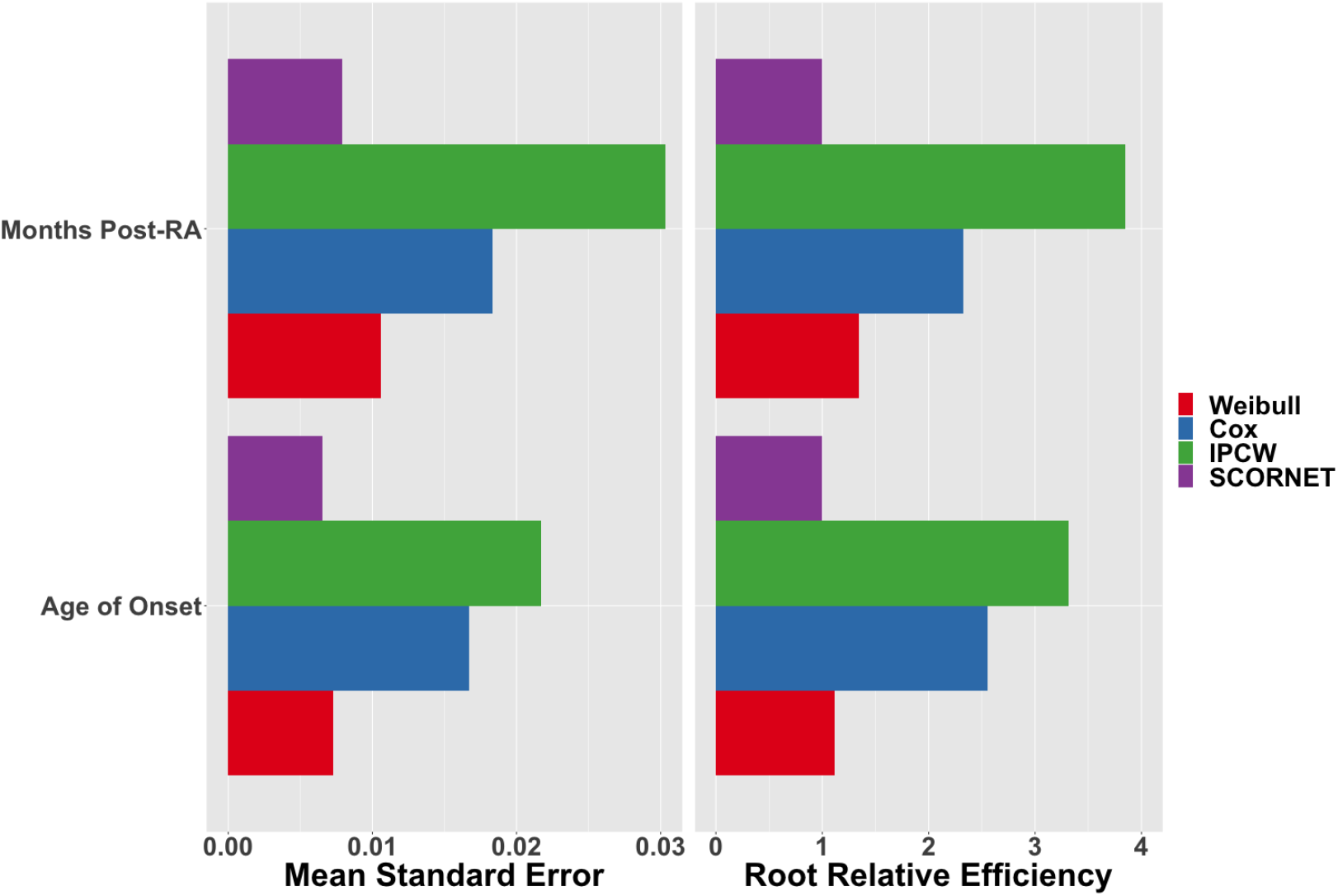
Time-averaged bootstrap standard errors (left) and empirical root relative efficiencies (right) of the Weibull Accelerated Failure Time (red), Cox Proportional Hazards w/ Breslow baseline (blue), IPCW (green), and SCORNET (purple) estimators for estimation of (1) age-specific HF risk (left), and (2) HF risk after RA diagnosis (right), among RA patients in the Partners EHR database.

## 5 Discussion

By leveraging a sizeable unlabeled data set containing imperfect surrogates of the true event times and a small set with observed current status labels, the SCORNET estimator serves as a robust and efficient alternative to existing model-free survival estimators with current status data. The semi-supervised nature of SCORNET makes it well-conditioned to EHR-based survival estimation in settings where only a limited number of labels are available or readily obtainable. Moreover, by only requiring current status labels rather than the precise timing of event onset, SCORNET greatly reduces the burden of chart review and increases the feasibility of studying disease risk using EHR data.

To allow for covariate-dependent censoring, which is frequently present in observational settings, SCORNET requires additional assumptions on the distribution of *C* | ***Z***_0_. Although we choose the proportional hazards model for illustration, any standard semi-parametric model will yield similar properties for the resulting estimator. Since {*C*, ***Z***_0_} are observed for all subjects, one can potentially allow for more flexible (i.e. non-parametric) censoring models. That said, our simulation results suggest that SCORNET is relatively insensitive to misspecification of the model for *C* | ***Z***_0_. Even under mild misspecification, it achieves consistently lower mean squared errors than existing estimators.

When interest lies in assessing how risk differs across different patient sub-populations, it is straightforward to extend SCORNET to estimate subgroup-specific risks for a small number of subgroups. However, future research is warranted to estimate covariate-specific risks for a general set of covariates.

## 6 Software

An R package, including a sample use case and complete documentation, is available at https://cran.r-project.org/web/packages/SCORNET/index.html. Source code can be found at https://github.com/celehs/SCORNET.

## Funding

This work was supported by the U.S. National Institutes of Health Grants T32-AR05588512, T32-GM7489714, and R21-CA242940.

## Acknowledgements

The authors declare no conflicts of interest.

## Notes

### Competing Interest Statement

The authors have declared no competing interest.

