## Supplemental Materials for "Semi-supervised Calibration of Risk with Noisy Event Times (SCORNET) Using Electronic Health Record Data"

### Supplementary Materials

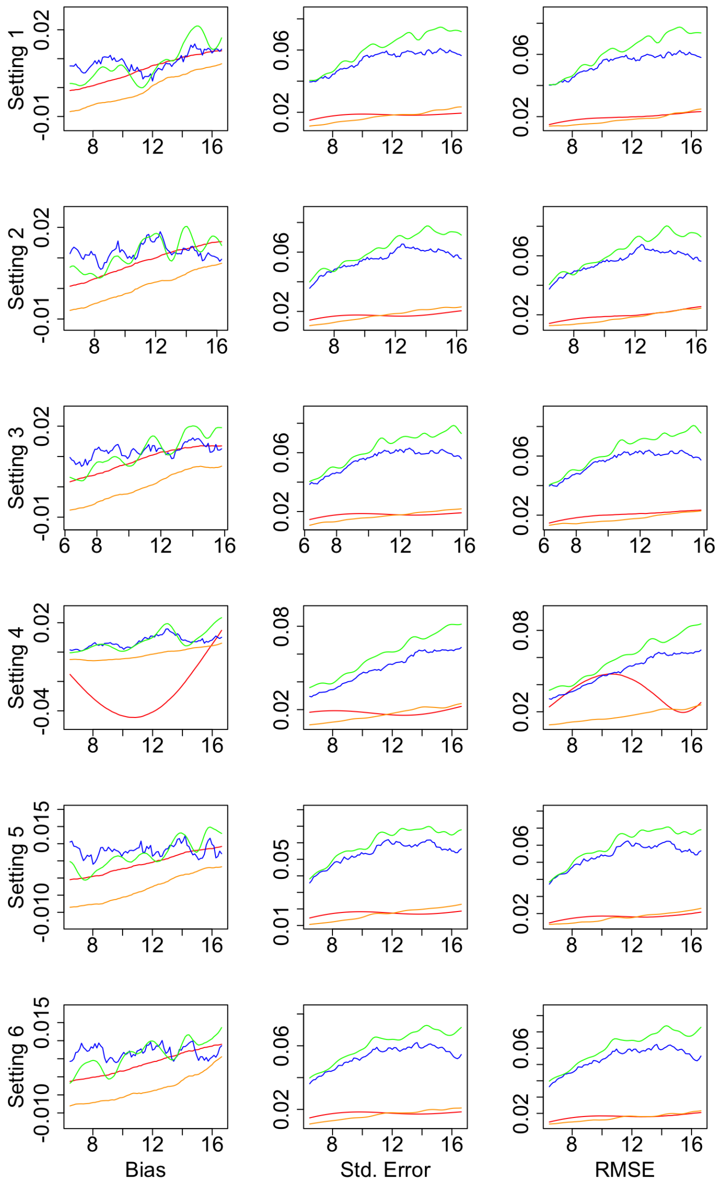


Figure 1: Empirical biases, standard errors, and root mean squared errors (RMSEs) over time of the Weibull Accelerated Failure Time (blue), Cox Proportional Hazards w/ Breslow baseline (red), supervised IPCW (green), and SCORNET-Strong (orange) estimators in various simulated settings with $n = 200$ observed current status labels. See Main Text Table 1 for details of the generative mechanism employed in each setting.

| | | $n = 100$ | | | | $n = 200$ | | | |
|---|---|---|---|---|---|---|---|---|---|
| Metric | Setting | Weibull | CoxPH | IPCW | SCORNET | Weibull | CoxPH | IPCW | SCORNET |
| \|Bias\| | 1 | .008 | .015 | .017 | .007 | .006 | .009 | .008 | .005 |
| | 2 | .009 | .015 | .018 | .006 | .009 | .012 | .012 | .004 |
| | 3 | .008 | .014 | .016 | .006 | .009 | .012 | .012 | .004 |
| | 4 | .029 | .009 | .013 | .006 | .029 | .007 | .008 | .004 |
| | 5 | .006 | .013 | .013 | .006 | .004 | .008 | .007 | .004 |
| | 6 | .006 | .012 | .013 | .006 | .004 | .007 | .006 | .004 |
| Emp SE | 1 | .026 | .068 | .088 | .021 | .018 | .054 | .061 | .017 |
| | 2 | .026 | .070 | .091 | .021 | .017 | .055 | .062 | .017 |
| | 3 | .026 | .068 | .089 | .021 | .018 | .055 | .063 | .017 |
| | 4 | .024 | .063 | .086 | .020 | .018 | .050 | .060 | .017 |
| | 5 | .025 | .068 | .087 | .021 | .017 | .054 | .060 | .017 |
| | 6 | .024 | .068 | .087 | .020 | .017 | .053 | .061 | .017 |
| RRE | 1 | 1.24 | 3.19 | 4.14 | 1.00 | 1.05 | 3.10 | 3.55 | 1.00 |
| | 2 | 1.22 | 3.29 | 4.29 | 1.00 | 1.00 | 3.17 | 3.60 | 1.00 |
| | 3 | 1.24 | 3.32 | 4.31 | 1.00 | 1.06 | 3.27 | 3.73 | 1.00 |
| | 4 | 1.23 | 3.20 | 4.38 | 1.00 | 1.09 | 3.01 | 3.60 | 1.00 |
| | 5 | 1.18 | 3.23 | 4.13 | 1.00 | 1.03 | 3.20 | 3.57 | 1.00 |
| | 6 | 1.19 | 3.31 | 4.26 | 1.00 | 1.06 | 3.22 | 3.66 | 1.00 |
| RRMSE | 1 | 1.23 | 3.06 | 3.97 | 1.00 | 1.09 | 3.00 | 3.43 | 1.00 |
| | 2 | 1.25 | 3.20 | 4.16 | 1.00 | 1.10 | 3.13 | 3.53 | 1.00 |
| | 3 | 1.25 | 3.21 | 4.16 | 1.00 | 1.15 | 3.21 | 3.65 | 1.00 |
| | 4 | 1.89 | 3.03 | 4.15 | 1.00 | 2.05 | 2.94 | 3.52 | 1.00 |
| | 5 | 1.16 | 3.10 | 3.95 | 1.00 | 1.02 | 3.08 | 3.42 | 1.00 |
| | 6 | 1.16 | 3.16 | 4.05 | 1.00 | 1.04 | 3.10 | 3.51 | 1.00 |

Table 1: Time-averaged empirical absolute biases and standard errors as well as root relative efficiencies and relative root mean squared errors (compared to SCORNET) of the Weibull Accelerated Failure Time, Cox Proportional Hazards w/ Breslow baseline, supervised IPCW, and SCORNET estimators on simulated datasets with $n = 100$ and $n = 200$ observed current status labels. See Main Text Table 1 for details of the generative mechanism employed in each simulation setting.
